# BioSamplr: An open source, low cost automated sampling system for bioreactors

**DOI:** 10.1101/2020.10.25.354183

**Authors:** John P. Efromson, Shuai Li, Michael D. Lynch

## Abstract

Autosampling from bioreactors reduces error, increases reproducibility and offers improved aseptic handling when compared to manual sampling. Additionally, autosampling greatly decreases the hands-on time required for a bioreactor experiment and enables sampling 24 hrs a day. We have designed, built and tested a low cost, open source, automated bioreactor sampling system, the BioSamplr. The BioSamplr can take up to ten samples from a bioreactor at a desired sample interval and cools them to a desired temperature. The device, assembled from low cost and 3D printed components, is controlled wirelessly by a Raspberry Pi, and records all sampling data to a log file. The cost and accessibility of the BioSamplr make it useful for laboratories without access to more expensive and complex autosampling systems.

## 1. Hardware in context

Fermentations must be monitored closely, sometimes over long periods of time. Consistent sampling, even at odd-hours, is essential and can be labor intensive. Automated sampling can reduce this burden, while additionally increasing sterility, removing user error and improving reproducibility. Autosampling is not a new concept, with one of the first systems for aseptic sampling described in 1987 by Seiffert and Matteau ^1^ Despite significant developments in bioreactor designs over the past 30 years, autosampling has still not become commonplace in most laboratories. Currently, while multiple commercial automated sampling systems do exist they are quite expensive, which has limited their adoption by many labs. The BioSamplr, is an accessible option, costing approximately $700, (minus the cost of the sampling probe) and assembled from affordable materials and 3D printed parts. All functions are run via a Raspberry Pi controller using open source software. The BioSamplr takes samples at desired time intervals and holds them at a target temperature. The current design enables up to 10 samples dispensed into chilled, standard 1.5mL microcentrifuge tubes. Sample cooling has been validated for up to 30 hours at 4 degrees Celsius. Fluid sensors validate a successful sampling event and all temperature and sampling data is recorded to log files. To our knowledge this is the first open source bioreactor sampling instrument.

As mentioned, there are several commercial autosampling systems available for bioreactors. These various systems offer advanced features and capabilities. For example, some can prepare pre-diluted samples while others can deliver aliquots to up to five different analytical instruments. Many designs are modular with multiple components including fraction collectors, valve manifolds, and sterile filtering probes. A few commercial autosampling systems available for lab bioreactors include the BioProbe™ from Bbi-Biotech, the Omnicoll Fraction Collector and Sampler from Lambda Instruments and the Segflow™ by Flownamics, and more general reactor sampling system, the EasySampler™ by Mettler Toledo. A summary of some of the key features of these systems with a comparison to the BioSamplr is given in Table 2 below.

**Table 1:** Specifications table.

| **Table 1: Specifications table** |  |
| --- | --- |
| **Hardware name** | BioSamplr |
| **Subject area** | ● Biological Sciences<br>● Bioengineering<br>● Fermentation |
| **Hardware type** | Biological sample handling and preparation |
| **Open Source License** | Attribution-ShareAlike 3.0 |
| **Cost of Hardware** | \$700 |
| **Source File Repositories** | Hardware: https://osf.io/r6jkg/<br>Software: https://github.com/DukeLynchLab/BioSamplr |

**Table 2:**
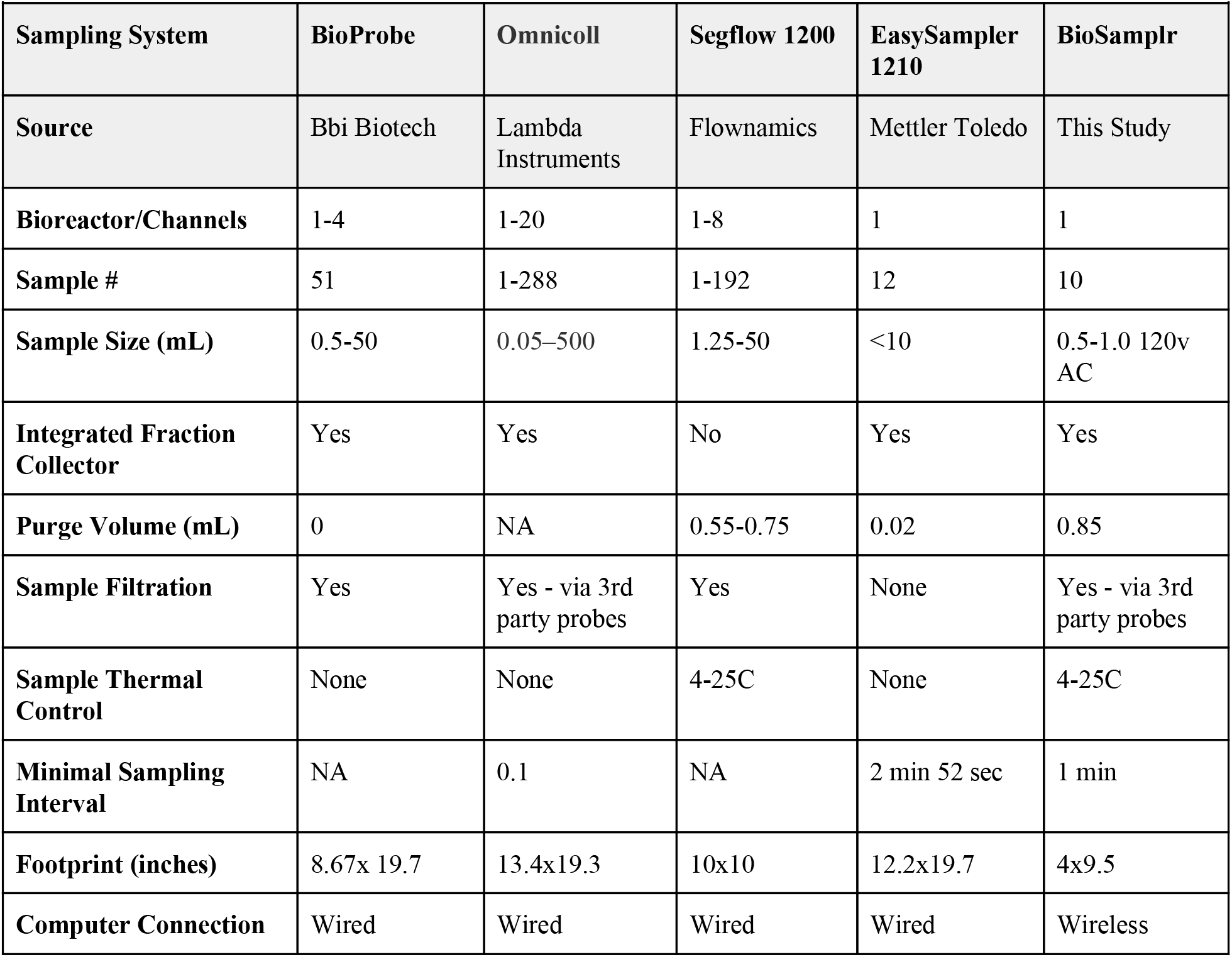
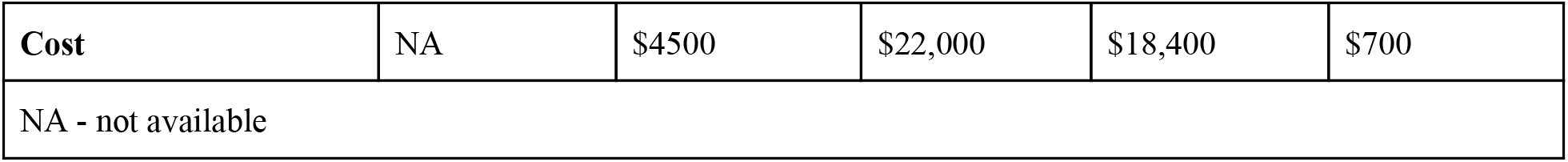
A comparison of bioreactor autosamplers.

## 2. Hardware description

The BioSamplr design was primarily based upon work by Cannizzaro and Von Stockar, (in a now abandoned US patent application) wherein sterile air can be used for “flushing of sample tubing between or during individual sampling steps”. ^2^ In this design (Figure 1a), a pump is used to pull the liquid/air and valves are used to switch from sampling the reactor to purging with filtered air. The BioSamplr hardware (Figure 1b-e) can be broken down into four primary systems i) the pumping/valve system, ii) the cartesian system, iii) the sample block, and v) the electronics.

**Figure 1:**
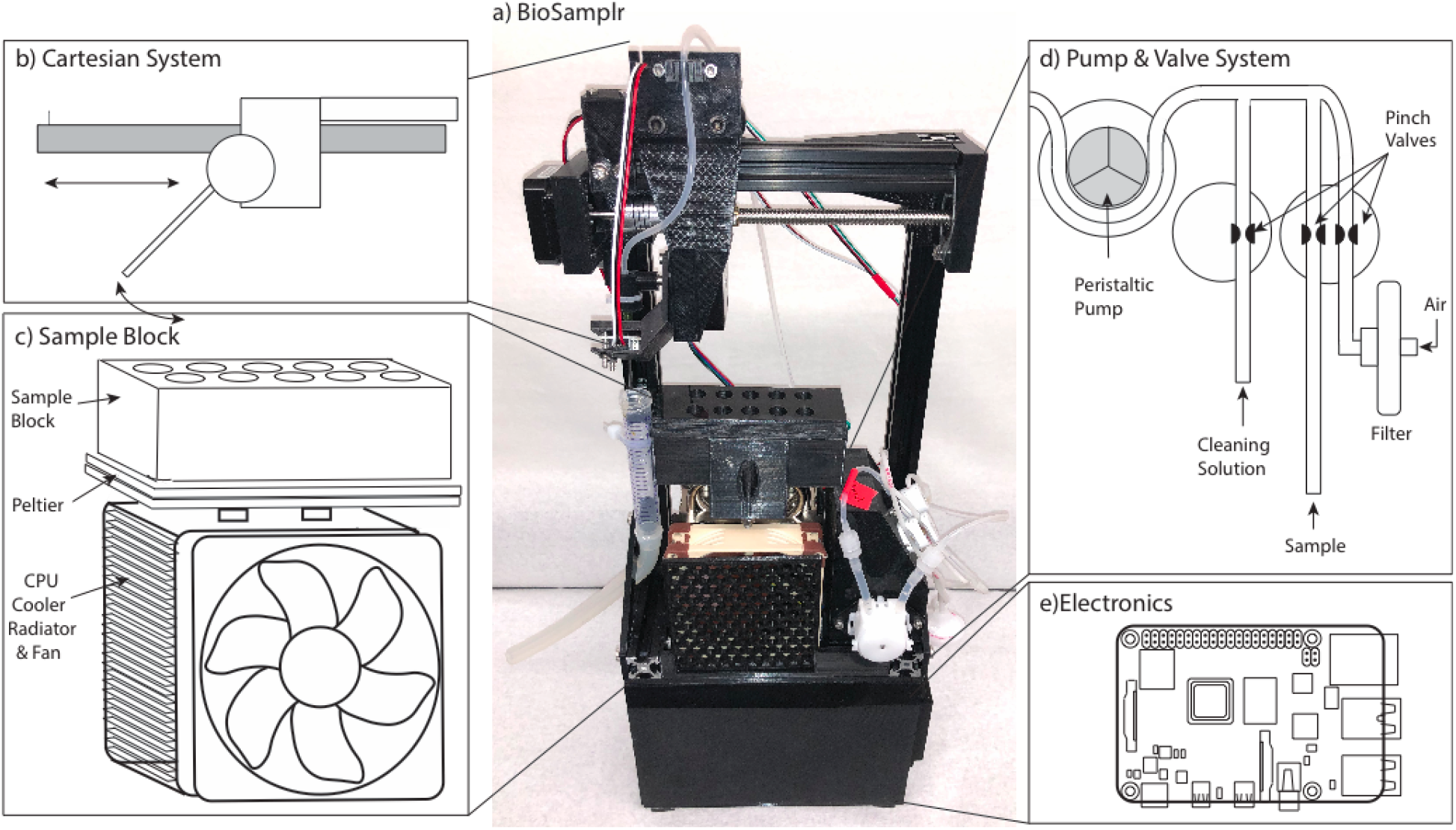
a) A picture of an assembled BioSamplr systems including: (b) the Cartesian System, which moves sampling needle to the appropriate sample on the (c) Sample Block, which holds and chills up to 10 microfuge tubes. Samples are drawn from a reactor by a (d) Pump and Valve System. This system uses a peristaltic pump to draw liquid (sample and or cleaning solution) or air into the system. Pinch valves control which medium is pumped. e) The electronics including a Raspberry Pi Controller are contained in a housing underneath the other components.

### 2.1 Cartesian System

The cartesian system (Figure 1b) locates the blunt sample dispensing needle over a target sample tube (or waste) based on x, y coordinates. The sample needle is moved to a given x,y coordinate by first moving a gantry (via an actuated lead screw) along an aluminum rail. The needle is centered between two rows of five sample tubes. A stepper motor is used to rotate the needle to dispense the sample in a given tube.

### 2.2 Sample Block

The sample block consists of a rack which holds tubes in a water filled, insulated aluminum box, cooled with peltiers, which in turn transfer heat to a CPU cooling unit containing a radiator and exhaust fan. Peltier modules are commonly used in cooling devices and more frequently in DIY thermal control equipment. ^3^ The temperature controlled sample block uses two custom fabricated thermal probes,^4^ to take readings from both the water bath and the radiator, which are then recorded and used to control the temperature. Pulse width modulation (PWM) is used to control the Peltier’s duty cycle turning it on and off in order to modulate temperature control. PWM has been compared to constant current circuitry and it has been shown that while energy efficiency decreases slightly, ^5^ the lifespan of the Peltier module is unaffected and the module can remain reliable over one hundred thousand hours. ^6^ With the current configuration the sample block has been validated for use up to 30 hours and can maintain a 4 degree Celsius setpoint within 1.2 degrees.

### 2.3 Pump/Valve System

The pumping system uses two 24V three-way pinch valves to direct fluid flow from the bioreactor, a cleaning solution, or external sterile filtered air. Liquid/air is pumped through sample tubing to a blunt sample dispensing needle. These fluids are propelled through the flowpath by a 12V DC peristaltic pump. Fluid sensors along the flow path confirm that a sample has actually been taken.

### 2.4 Electronics Housing

The electronics, two power source modules (24V, 6-9 A and 5V, 3A) and the Raspberry Pi controller are contained in a 3D printed housing beneath the cartesian system. Each system is discussed below. Real time cell free sampling can be performed using commercially available probes, such as the ceramic FISP probe (obtainable from Flownamics (Amsterdam, Netherlands) or from YSI (Yellow Springs, OH SKU 002850) or the BioProbe Sampling probe (obtainable from bbi Biotech, Berlin, Germany).

## 3. Design Files

### 3.1 3D Print Files

There are twenty four different 3D print files, some of which include multiple pieces. These include all BioSamplr components that are not directly purchased. All design/print files are obtainable at https://osf.io/r6jkg/.

- The “Needle Holder” attaches a female luer-lok connector with an 18 gauge blunt sample dispensing needle to the y-axis stepper motor.
- “M5_7mm_Spacer.stl” includes four 7mm spacers that are used to space the wheels of the X Gantry from the Ganty. Only two are necessary but as they are a small part, extra are included.
- “Part 1 X Gantry” is the main section of the gantry which holds the y-axis stepper motor and connects to Part 2 of the X Gantry. This gantry also holds the y-axis endstop switch.
- “Part 2 X Gantry” connects Part 1 of the X Gantry to wheels which mount on the x-axis rail. This piece also holds the second fluid sensor.
- The “Pillow Bracket” holds the lead screw pillow which supports one end of the lead screw and connects to the right leg of the cartesian system outside of the right top bracket.
- The “Radiator Bracket” holds the radiator and connects the two legs together.
- The “Sample Block Lid” holds ten microcentrifuge sample tubes in the sample block.
- The “Sample Block” is a hollow walled box which holds an aluminum box and connects to the radiator holding the Peltier in place as well as the aluminum box to the Peltier. The hollow walls can be filled with spray foam insulation.
- The “Screw Stepper Bracket” mounts the x-axis stepper motor to the left leg of the cartesian system.
- The “Top Brackets” connect the x-axis aluminum rail to the two legs of the cartesian system.
- The “Waste Tube Bracket” attaches to the left leg of the cartesian system and has a notch cutout on the backside for a zip-tie. This zip-tip will hold a 15mL centrifuge tube with a cut off bottom that is used as a funnel to direct waste liquids into the waste tubing and further to the waste location.
- The “Wire Clips” mount into the v-slot of the aluminum extrusion rails and can tether wires to the rails for cable management. They are meant for the back side of the cartesian system and while only four are necessary they are a small part, so twelve are included.
- The “X Switch Bracket” mounts an endstop switch to the leftmost side of the x-axis aluminum rail. The X Gantry triggers this switch to signal its location on the x-axis.
- The “Y Switch Bracket” mounts an endstop switch to the X-Gantry. The sample dispensing needle triggers this switch to signal its location rotating on the y-axis.
- The two “YZ Brackets” connect the legs of the cartesian system to the two aluminum rails and the Radiator Bracket.
- The “Fan Wall” is part of the circuitry housing and mounts a system cooling fan using the System Fan Bracket. This wall connects to the Housing Bottom, Housing Top, Power Source Wall and Pi Wall.
- The “Housing Bottom” is part of the circuitry housing and mounts the main PCB using heat set inserts while also connecting to other walls of the circuitry housing. This surface connects to all walls of the circuitry housing.
- The “Housing Top” is part of the circuitry housing and mounts the cartesian system to the circuitry housing using heat set inserts. This surface connects to the Pi Wall, Fan Wall, Vent Wall, and the base aluminum rails of the cartesian system.
- The “Pi Wall” is the back wall of the circuitry housing and has locations to mount a Raspberry Pi as well as a dual channel relay board by way of heat set inserts. This wall connects to the Housing Bottom, Housing Top. Fan Wall and Vent Wall.
- The “Power Source Wall” is the front wall of the circuitry housing and has locations for the two buck power supply modules as well as a step-down buck converter module all using heat set inserts. This wall connects to the Housing Bottom, Fan Wall, and Vent Wall.
- The “Safety Bracket” mounts over the power socket wiring on the inside of the circuitry housing to provide some safety from live wires.
- The “System Fan Bracket” mounts to the Fan Wall and holds the system cooling fan in place.
- The “Vent Wall” has a socket for the main power socket as well as vents to allow aeration of electronics. This wall connects to the Housing Bottom, Housing Top, Pi Wall and Power Source Wall.

### 3.2 Electronics

There are two custom electronic systems, the power source and controller PCB board, which are integrated with other electronic components. These are described in the three files below.

- The “BioSamplr Gerber” file is the layout of the main PCB that controls the electronics of the sampler.
- The “BioSamplr PCB Schematic” outlines all connections within the circuitry of the device.
- The “BioSamplr Powersource Schematic” outlines the connections from the power socket powersource to the buck powersource modules and the buck step down converter with terminal connections to the PCB and Raspberry Pi by way of a DC barrel adapter.

### 3.3 Software

As mentioned above, the BioSamplr is controlled via a Raspberry Pi and python scripts. The scripts can be divided into 2 categories, those used for system initialization and calibration and those used for operation. These scripts are described below.

#### Initialization/Calibration Scripts

- “boot.py” is a boot file that when added to the Raspberry Pi startup process, sets both valves to their initial positions.
- “GPIO_cleanup.py” is a script that initializes all GPIO pins of use on the Raspberry Pi and sets them all to “LOW” which turns off the pump, and Peltier while setting the valves to their default position.
- “Cartesian_Test.py” is a script that locates the sample dispensing needle over each sample tube to test location coordinates
- “Photo_Transistor_Test.py” is a script that reads the two analog to digital converter channels for the two fluid sensors and converts their raw values into a value between 0 and 1. This script is used for calibrating fluid sensors.
- “Pump_Test.py” is a script that has all pump and valve functions so that they can easily be controlled one by one. The purpose of this script is to validate that the pump and valves are functioning properly.
- “sleep.py” is a script that activates the stepper motors for five minutes to allow a voltage reading to be measured from the motor driver chips on the main PCB. This is necessary during setup to calibrate the current going to the stepper motors
- “simpletest.py” is a script written by Tony DiCola Adafruit Industries ^7^. This script reads the eight different channels of the analog to digital converter and prints their values to the terminal. The purpose of this script is to validate that the thermal probes and fluid sensors are reading properly as well as that the other four channels are properly grounded and reading zero.

#### Operation Scripts

- “BioSamplr_Master.py” is the master control python script needed to control sampling. This control script will take ten samples at a user input time-interval and keep the sample block at a setpoint temperature while logging sampling data to a log file..
- “autoHome.py” is a script that first rotates the sample dispensing needle on the y-axis until it triggers the y-axis endstop switch, then rotates back to the vertical position between the two rows of sample tubes. Secondly it moves the x gantry along the x axis until it engages the x-axis endstop switch and then backs off this switch slightly. The purpose of this motion is to “home” the sample dispensing needle over a reference location that is also the location of the waste tube.
- “Log_Temp.py” is a script that activates the sample block temperature control and logs temperature readings from two thermal probes to a log file.
- “Measure_Temp.py” is a script that reads two thermal probes and prints their resulting temperature values to the terminal.
- “Cleaning_Cycle.py” is a cleaning cycle script that gives instructions to place sample tubing in a cleaning solution or air to clean sample tubing.

## 4. Bill of Materials

The bill of materials can be found at https://osf.io/r72d6/

## 5. Build Instructions & Operation

Due to the complexity of the instrument, we have prepared several videos which walk through the detailed BioSamplr assembly, calibration and unit operation including software set up. Links to these videos are found in Table 4 below. With a single low speed 3D printer, printing of components can be expected to take about 5 days. For someone with experience with these type of DIY projects as well as Raspberry Pi, ~3 days should be planned for assembly and calibration. Additionally, to fabricate two thermal probes, follow the directions found online to make a waterproof negative temperature coefficient (NTC) thermistor probe. ^4^

**Table 3:**
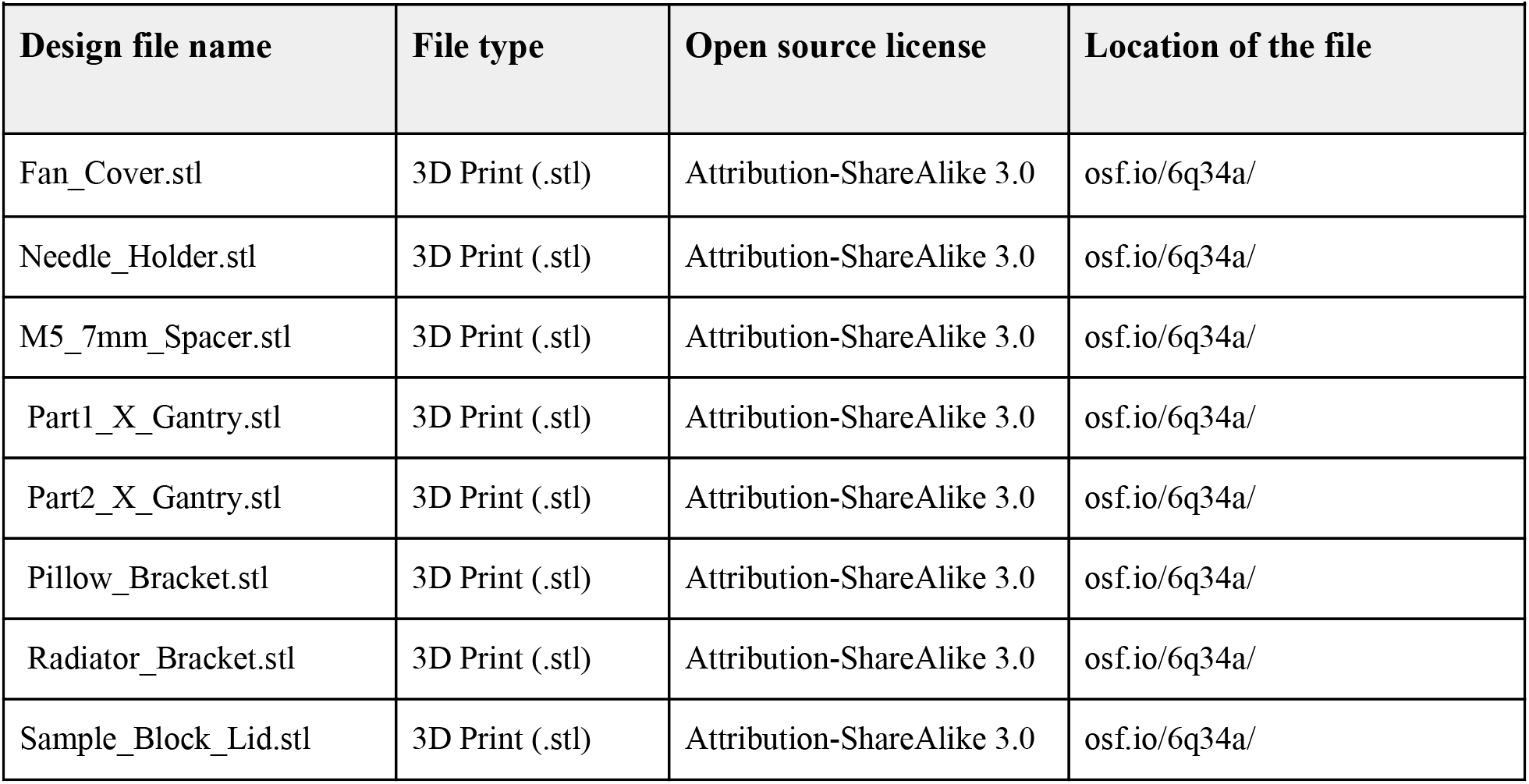
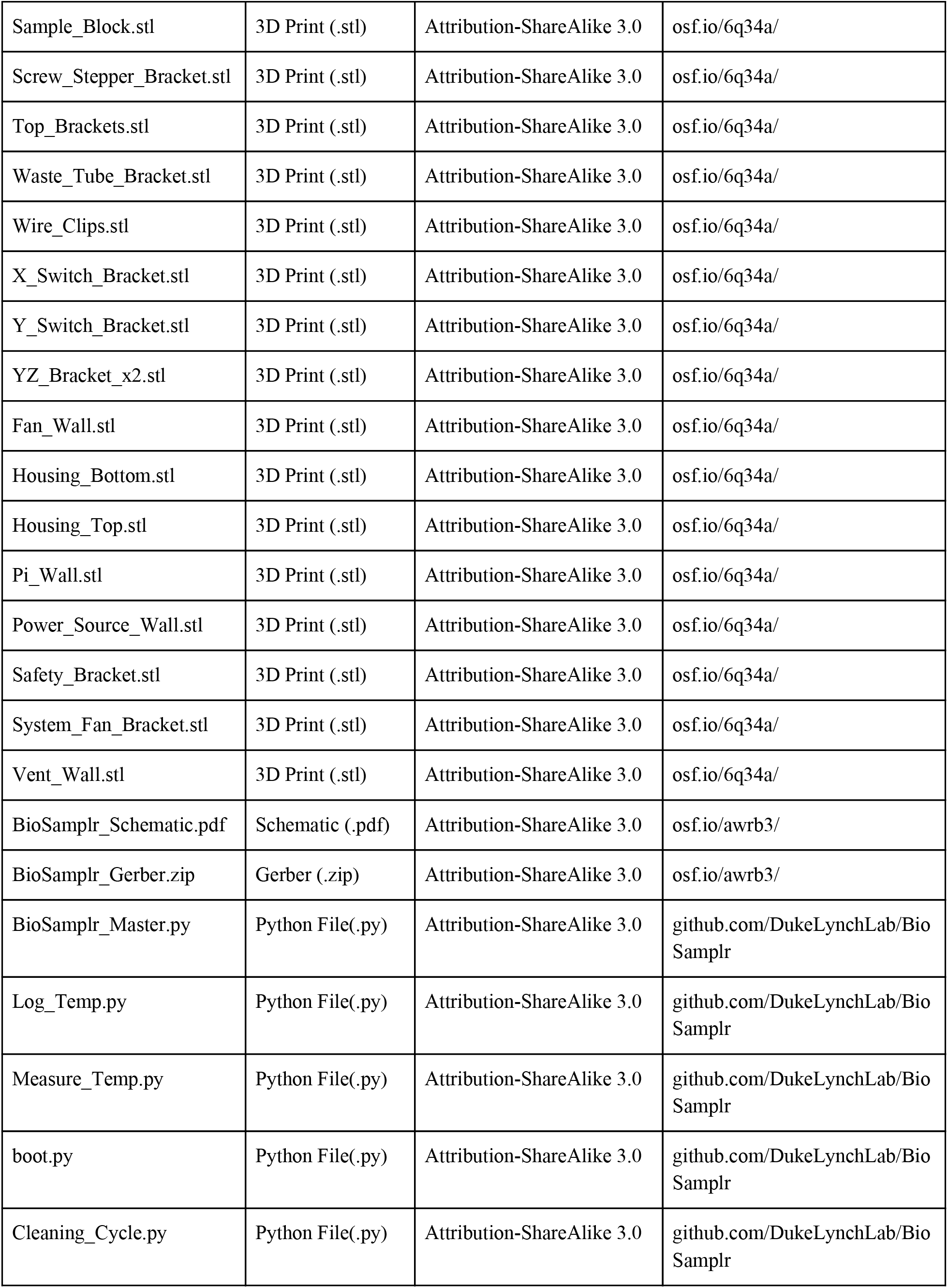
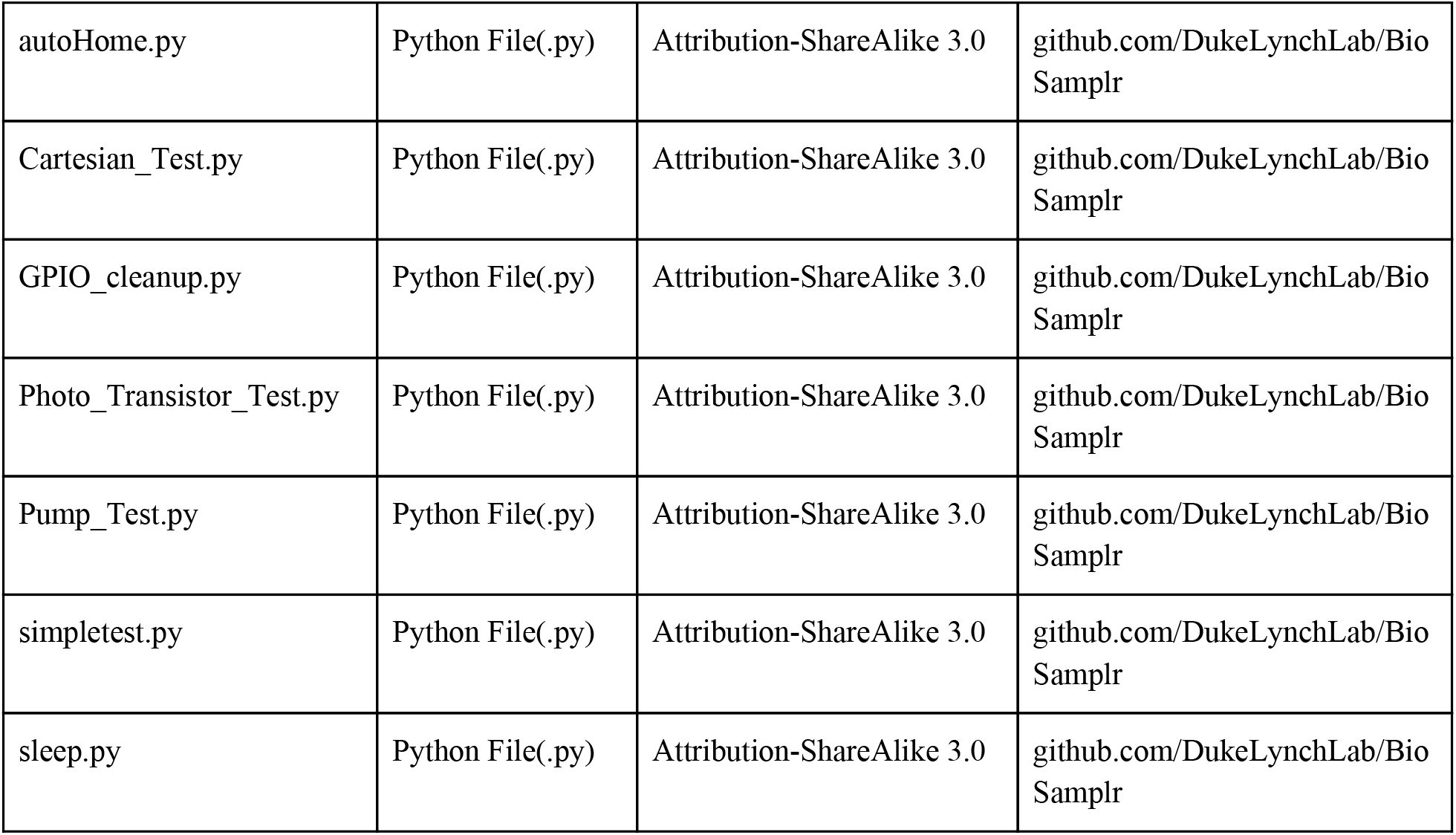
Design File Summary.

| **Table 3: Design File Summary** |  |  |  |
| --- | --- | --- | --- |
| **Design file name** | **File type** | **Open source license** | **Location of the file** |
| Fan_Cover.stl | 3D Print (.stl) | Attribution-ShareAlike 3.0 | osf.io/6q34a/ |
| Needle_Holder.stl | 3D Print (.stl) | Attribution-ShareAlike 3.0 | osf.io/6q34a/ |
| M5_7mm_Spacer.stl | 3D Print (.stl) | Attribution-ShareAlike 3.0 | osf.io/6q34a/ |
| Part1_X_Gantry.stl | 3D Print (.stl) | Attribution-ShareAlike 3.0 | osf.io/6q34a/ |
| Part2_X_Gantry.stl | 3D Print (.stl) | Attribution-ShareAlike 3.0 | osf.io/6q34a/ |
| Pillow_Bracket.stl | 3D Print (.stl) | Attribution-ShareAlike 3.0 | osf.io/6q34a/ |
| Radiator_Bracket.stl | 3D Print (.stl) | Attribution-ShareAlike 3.0 | osf.io/6q34a/ |
| Sample_Block_Lid.stl | 3D Print (.stl) | Attribution-ShareAlike 3.0 | osf.io/6q34a/ |
| Sample_Block.stl | 3D Print (.stl) | Attribution-ShareAlike 3.0 | osf.io/6q34a/ |
| Screw_Stepper_Bracket.stl | 3D Print (.stl) | Attribution-ShareAlike 3.0 | osf.io/6q34a/ |
| Top_Brackets.stl | 3D Print (.stl) | Attribution-ShareAlike 3.0 | osf.io/6q34a/ |
| Waste_Tube_Bracket.stl | 3D Print (.stl) | Attribution-ShareAlike 3.0 | osf.io/6q34a/ |
| Wire_Clips.stl | 3D Print (.stl) | Attribution-ShareAlike 3.0 | osf.io/6q34a/ |
| X_Switch_Bracket.stl | 3D Print (.stl) | Attribution-ShareAlike 3.0 | osf.io/6q34a/ |
| Y_Switch_Bracket.stl | 3D Print (.stl) | Attribution-ShareAlike 3.0 | osf.io/6q34a/ |
| YZ_Bracket_x2.stl | 3D Print (.stl) | Attribution-ShareAlike 3.0 | osf.io/6q34a/ |
| Fan_Wall.stl | 3D Print (.stl) | Attribution-ShareAlike 3.0 | osf.io/6q34a/ |
| Housing_Bottom.stl | 3D Print (.stl) | Attribution-ShareAlike 3.0 | osf.io/6q34a/ |
| Housing_Top.stl | 3D Print (.stl) | Attribution-ShareAlike 3.0 | osf.io/6q34a/ |
| Pi_Wall.stl | 3D Print (.stl) | Attribution-ShareAlike 3.0 | osf.io/6q34a/ |
| Power_Source_Wall.stl | 3D Print (.stl) | Attribution-ShareAlike 3.0 | osf.io/6q34a/ |
| Safety_Bracket.stl | 3D Print (.stl) | Attribution-ShareAlike 3.0 | osf.io/6q34a/ |
| System_Fan_Bracket.stl | 3D Print (.stl) | Attribution-ShareAlike 3.0 | osf.io/6q34a/ |
| Vent_Wall.stl | 3D Print (.stl) | Attribution-ShareAlike 3.0 | osf.io/6q34a/ |
| BioSamplr_Schematic.pdf | Schematic (.pdf) | Attribution-ShareAlike 3.0 | osf.io/awrb3/ |
| BioSamplr_Gerber.zip | Gerber (.zip) | Attribution-ShareAlike 3.0 | osf.io/awrb3/ |
| BioSamplr_Master.py | Python File(.py) | Attribution-ShareAlike 3.0 | github.com/DukeLynchLab/BioSamplr |
| Log_Temp.py | Python File(.py) | Attribution-ShareAlike 3.0 | github.com/DukeLynchLab/BioSamplr |
| Measure_Temp.py | Python File(.py) | Attribution-ShareAlike 3.0 | github.com/DukeLynchLab/BioSamplr |
| boot.py | Python File(.py) | Attribution-ShareAlike 3.0 | github.com/DukeLynchLab/BioSamplr |
| Cleaning_Cycle.py | Python File(.py) | Attribution-ShareAlike 3.0 | github.com/DukeLynchLab/BioSamplr |
| autoHome.py | Python File(.py) | Attribution-ShareAlike 3.0 | github.com/DukeLynchLab/BioSamplr |
| Cartesian_Test.py | Python File(.py) | Attribution-ShareAlike 3.0 | github.com/DukeLynchLab/BioSamplr |
| GPIO_cleanup.py | Python File(.py) | Attribution-ShareAlike 3.0 | github.com/DukeLynchLab/BioSamplr |
| Photo_Transistor_Test.py | Python File(.py) | Attribution-ShareAlike 3.0 | github.com/DukeLynchLab/BioSamplr |
| Pump_Test.py | Python File(.py) | Attribution-ShareAlike 3.0 | github.com/DukeLynchLab/BioSamplr |
| simpletest.py | Python File(.py) | Attribution-ShareAlike 3.0 | github.com/DukeLynchLab/BioSamplr |
| sleep.py | Python File(.py) | Attribution-ShareAlike 3.0 | github.com/DukeLynchLab/BioSamplr |

**Table 4:**
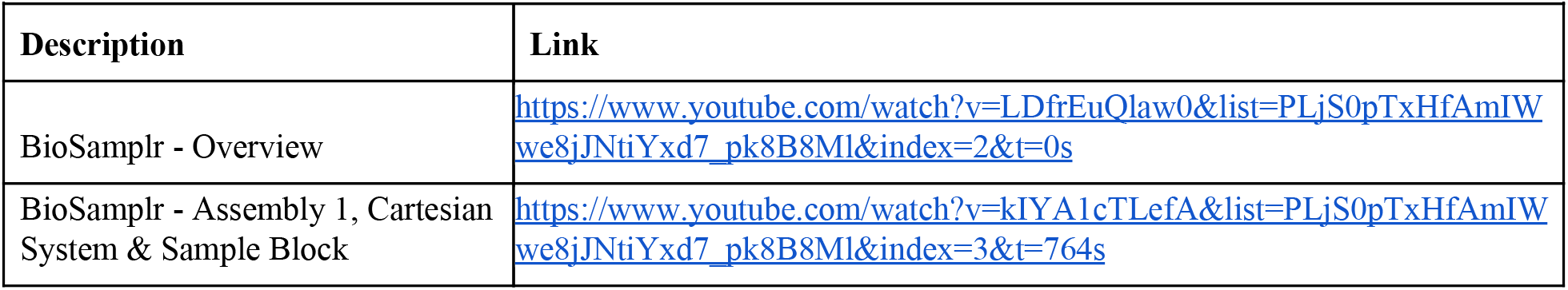
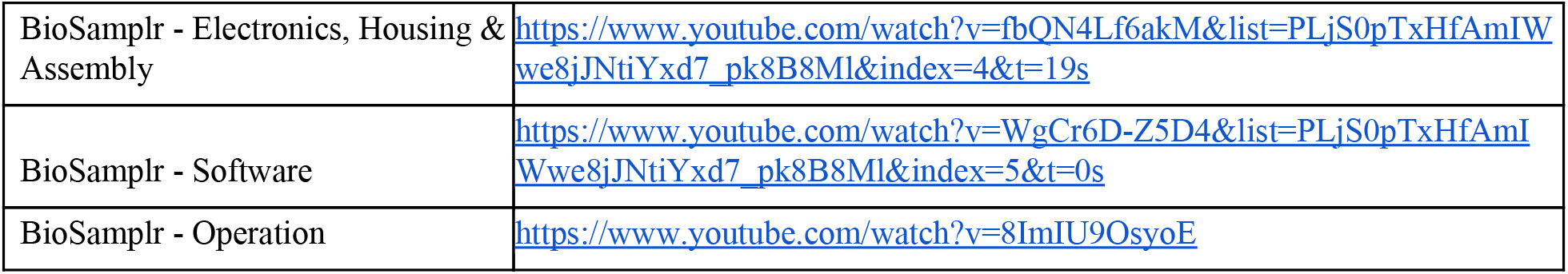
Instructional Videos.

### 5.1 3D printing

To begin, print all 3D printed components using the design files. There are twenty four different 3D print files, some of which include multiple pieces. These parts are listed in Table 5. All prints were performed using an Ender 3 Pro 3D Printer (Bill of Materials-Tools) with a layer height of 0.2mm, an infill density of 50%, a print temperature of 210 °C, a bed temperature of 60 °C, and a print speed of 50 mm/sec. We expect that any 3D printer can be used.

**Table 5:**
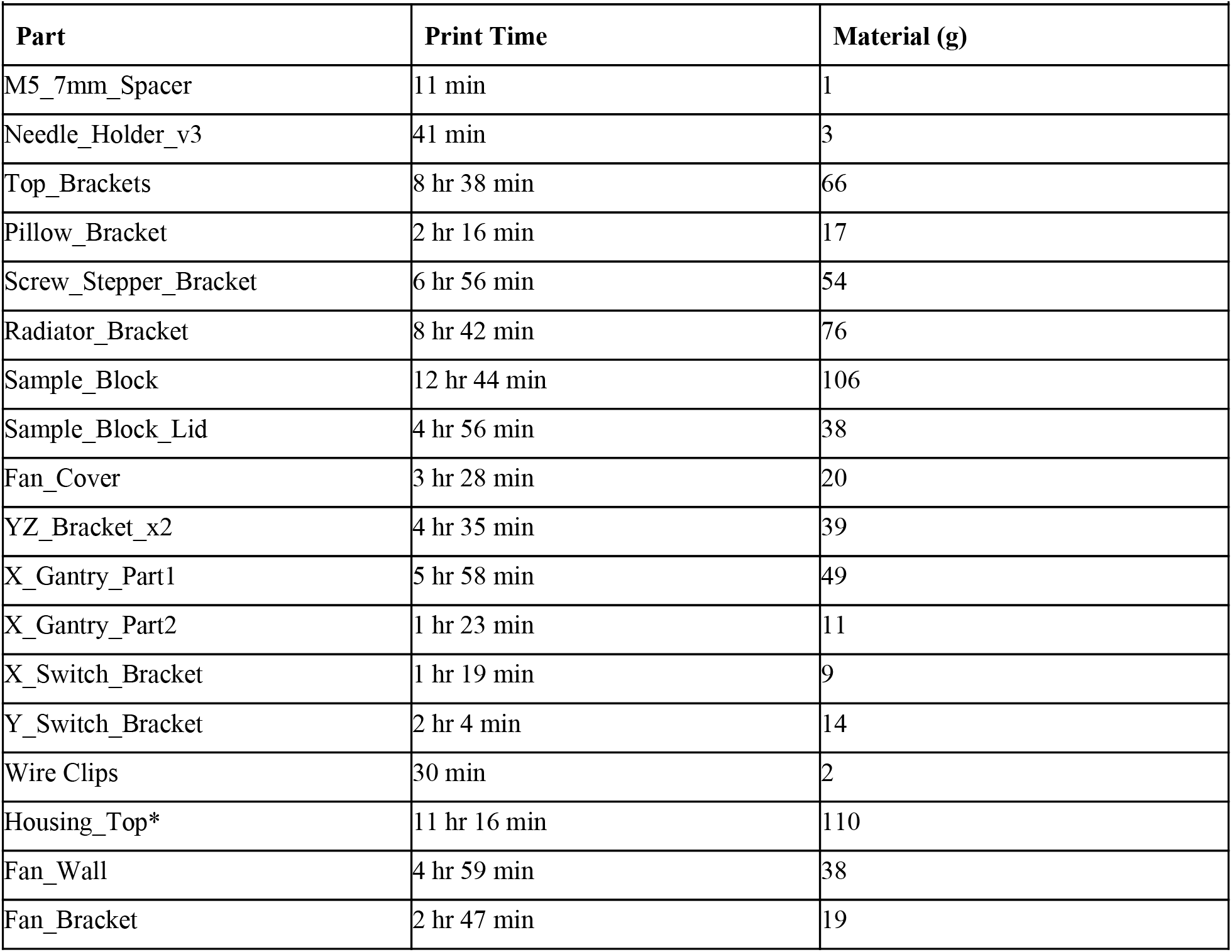
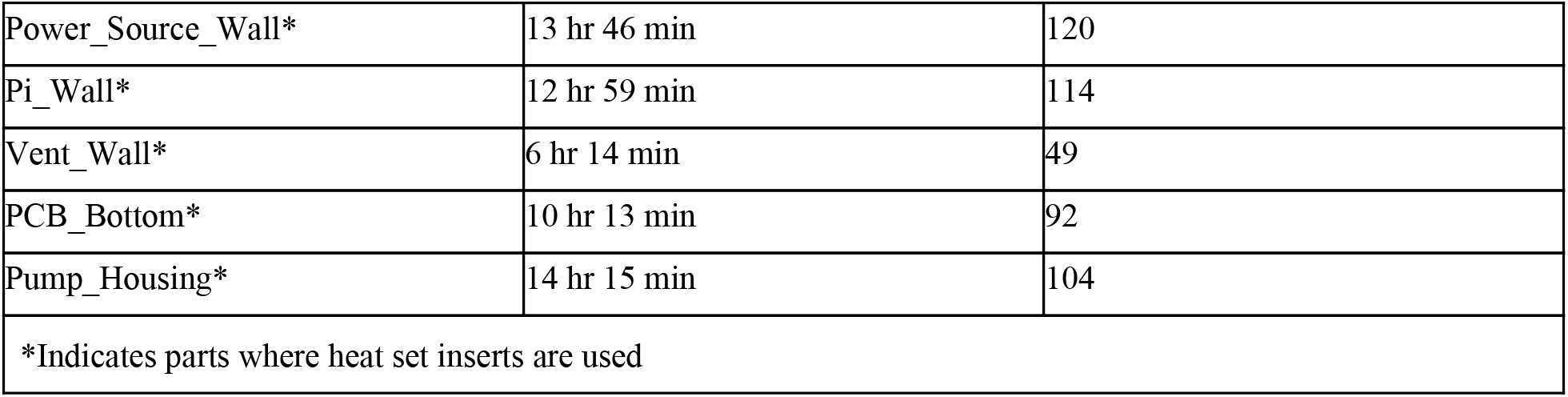
3D Printed Parts.

## 6. Instrument Operation

### 6.1 Device Safety

In addition to the details in the video, it is worth reemphasizing device safety. When plugged to a wall socket as intended, there are live 120V AC wires connecting the power socket to the two power modules as well as any exposed wiring inside the base housing. A safety bracket is installed covering the socket and the high voltage wires however fingers should be kept outside of the box while it is plugged in and live. The water in the sample block as well as the sample dispensing need and sample tubes are located above the electronics. While the electronics housing is water resistant it is not water-proof. Care should be taken to ensure that excessive liquid does not spill into the lower housing which could damage the instrument. The sample block should only be filled halfway with water or a water/bleach mixture as the sample tube’s volume will fill the remainder of the aluminum box.The sample dispensing needle moves along its trajectories as needed and is not covered or protected. Injury from this needle can be easily avoided following several precautions including using a blunt 18 gauge needle, keeping hands away from the needle while the device is live and/or operational and affixing it to the needle holder using a zip tie.

## 7. Validation and Characterization

### 7.1 Temperature Control

We first sought to validate the temperature control of the sample block. This was performed by monitoring the temperature of both the sample bath and radiator over, results of which are given in Figure 2 below. When a 4°C setpoint was used, the sample block was maintained within 1.2°C of the set point.

**Figure 2:**
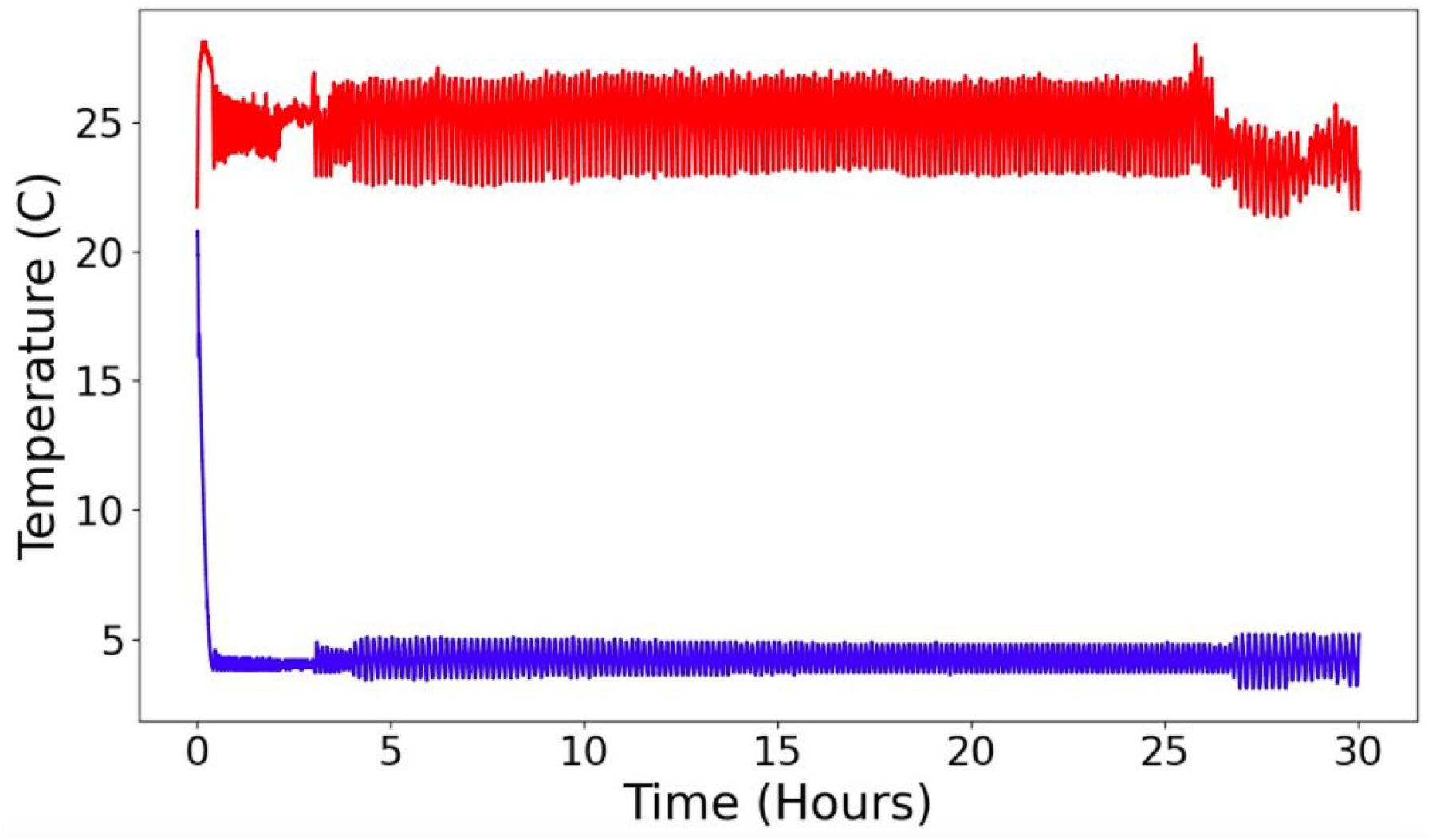
Temperature profile over time of the sample block (blue line) and radiator (red) line. Temperatures were monitored over 30 hours with a sample block temperature setpoint of 4°C setpoint.

### 7.2 Sample Volume Stability

In order to test the amount of evaporation from sample tubes the BioSamplr was set up to cool the sample block for a 30 hour time period at a 4 C temperature setpoint with ten sample tubes holding 1.5mL of water each. Initial weights of these tubes were recorded. Seven of these tubes were left open to the air while three were snapped shut. Following the designated time period the tubes were weighed again and these measurements were compared to the initial measurements. A difference of less than 1% (0.95%+/− 0.77%, n=10) from the initial measurements was observed.

### 7.3 Sample Volume Precision

Sample volume is a user defined variable. For purposes in our lab we routinely manually sample 1-5 mL from bioreactors. The peristaltic pump, while reliable, is not extremely accurate or precise, which does lead to some variation in sample volume. Other factors can influence this as well including variable sample viscosity throughout a fermentation process and bubbles that get into the sample tubing. In order to test variability of sample size the BioSamplr was set up to pump sixty samples of water (6 runs of 10 samples each) each targeting 0.75mL. The weight of each sample was measured. Results are given in Figure 3. The overall relative standard deviation was 25.7%., with the majority of samples within 0.2 mL of the target. 10% of the samples (6/60) were below 0.3mL less than half of the target. As having enough samples is critical for subsequent analysis, based on these results we would recommend target samples of > 1.0mL.

**Figure 3:**
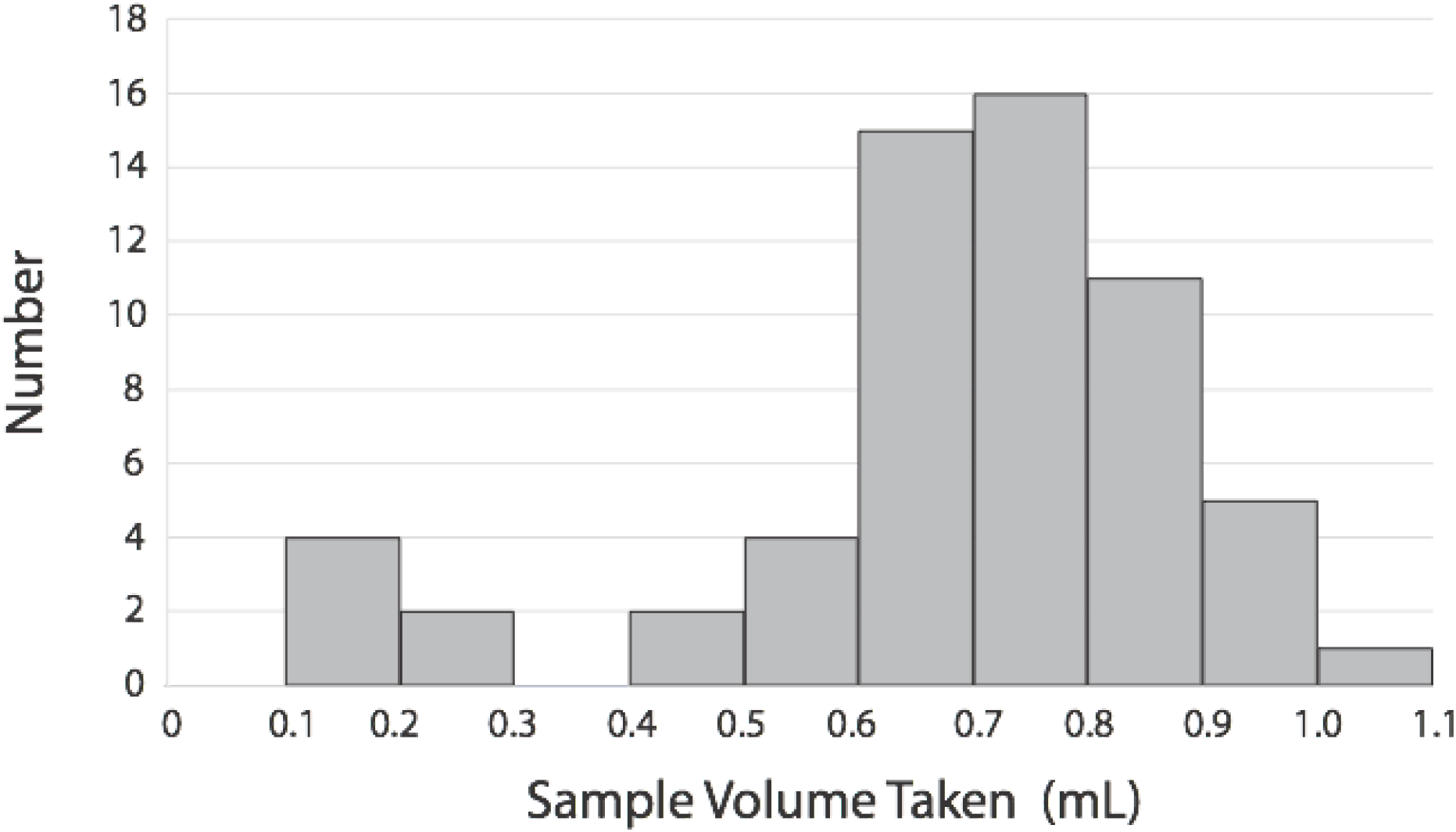
Sampling consistency for 6 runs of 10 samples (60 samples in total) all set to sample 0.75mL.

### 7.4 Fermentation Sampling

A fermentation using a genetically modified strain of *E. coli* producing pyruvate as recently reported by Li et al, ^8^ was repeated in this work, where in addition to manual sampling, the BioSamplr was used to take online/automated samples for a head to head comparison. Specifically, in this validation study we used a FISP™ (Flownamics, Amsterdam, Netherlands) cell free sampling probe to draw cell free samples from the bioreactor. Specifically, strain DLF_Z0043, with plasmid pCASCADE-g2, was used in two-stage fermentations, wherein phosphate depletion triggers a productive stationary phase and pyruvate production. This strain was evaluated in a 1L instrumented bioreactor (Infors Multifors), and two analytes (glucose and pyruvate) were measured from both manual and BioSamplr samples using HPLC as previously described. ^8^) Results are given in Figure 4 below. In the case of glucose the average relative error for automated samples compared to manual samples was 6±4% with a maximal error of 12%. In the case of glucose the average relative error for automated samples compared to manual samples was 2±2.7% with a maximal error of 8%.

**Figure 4:**
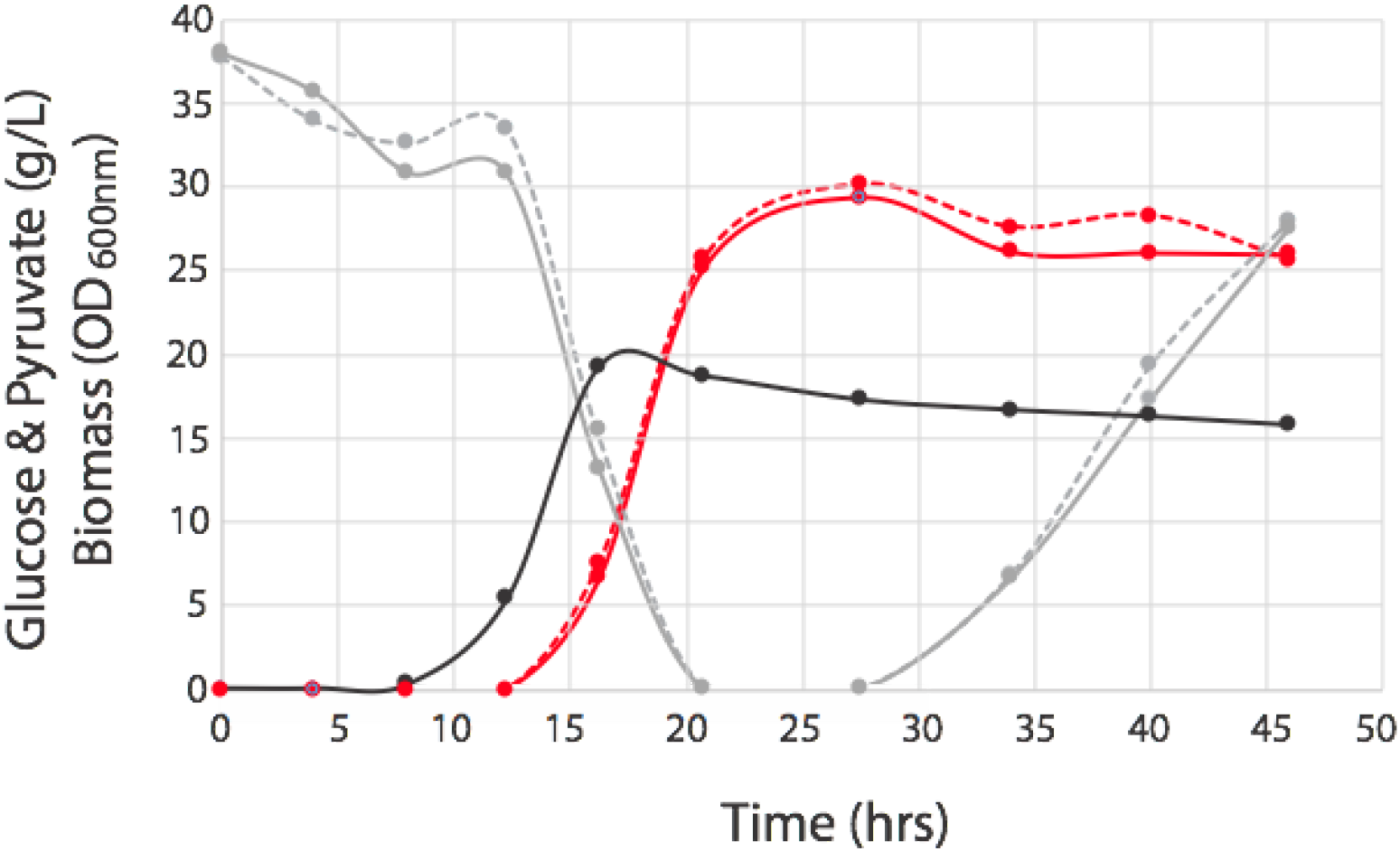
Comparison of data obtained from manual samples with those taken by the BioSamplr in actual fermentations. Cell free samples were taken and stored at 4°C. Optical density (black circles, manual samples), glucose measurements (gray circles and lines), pyruvate measurements (red circles and lines). Dashed lines indicate data from manual samples, whereas solid lines indicate samples taken with the BioSamplr.

## 8.0 Discussion

The combination of low cost microcontrollers and CPUs (such as Arduino and Raspberry Pi) and 3D printing has made DIY hardware for scientific applications more accessible than ever.^9^ Advanced hardware and control systems have been developed in numerous areas including in bioreactor control and data collection. ^10^ Commercial online probes/sensors are available for pO2, redox, pH, and temperature are routinely available, with probes reading optical density and/or viable cells becoming more commonplace. While advanced sensors, such as those based on raman spectroscopy, may be able to monitor metabolite and product levels in realtime, ^11,12^, sampling is still the gold standard used to measure analytes in fermentation broth. Manual sampling, particularly at regular intervals, is very time intensive and to date autosampling systems, which can significantly reduce the time required to execute a fermentation, have been expensive and inaccessible to many labs.

While the capabilities of commercial autosampling devices are useful (such as measuring flow rates ^13,14^) and perhaps essential in various contexts, they are not necessary for many, particularly those in an academic setting, where the primary value of automated sampling is reducing requirements for manual attention or sampling at odd hours. The BioSamplr, is an open source, low costs solution to autosampling. The initial version enables routine sampling, and refrigerated sample storage, greatly reducing the hands on time associated with bioreactor runs. Future iterations of the BioSamplr, will be focused on enabling larger sample numbers to extend unmanned operation, and perhaps the development of DIY sample probes.

## Acknowledgements

We would like to acknowledge support from DOE EERE grant #EE0007563 and DMC Biotechnologies, Inc.

## Author contributions

S. Li and J.P. Efromson tested the instrument. J.P. Efromson constructed the hardware and code. M.D. Lynch contributed to system design and analyzed results. All authors wrote, revised and edited the manuscript.

